# Automated Clear Cell Renal Carcinoma Grade Classification with Prognostic Significance

**DOI:** 10.1101/661520

**Authors:** Katherine Tian, Christopher A. Rubadue, Douglas I. Lin, Mitko Veta, Michael E. Pyle, Humayun Irshad, Yujing J. Heng

**Author notes:** Both authors contributed equally to this work (Katherine Tian, Christopher A. Rubadue). Corresponding Author: Dr. Jan Heng, Department of Pathology, Beth Israel Deaconess Medical Center, Harvard Medical School, 330 Brookline Ave, Dana 517B, Boston, MA 02215, USA.

## Abstract

We developed an automated 2-tiered Fuhrman’s grading system for clear cell renal cell carcinoma (ccRCC). Whole slide images (WSI) and clinical data were retrieved for 395 The Cancer Genome Atlas (TCGA) ccRCC cases. Pathologist 1 reviewed and selected regions of interests (ROIs). Nuclear segmentation was performed. Quantitative morphological, intensity, and texture features (*n*=72) were extracted. Features associated with grade were identified by constructing a Lasso model using data from cases with concordant 2-tiered Fuhrman’s grades between TCGA and Pathologist 1 (training set *n*=235; held-out test set *n*=42). Discordant cases (*n*=118) were additionally reviewed by Pathologist 2. Cox proportional hazard model evaluated the prognostic efficacy of the predicted grades in an extended test set which was created by combining the test set and discordant cases (*n*=160). The Lasso model consisted of 26 features and predicted grade with 84.6% sensitivity and 81.3% specificity in the test set. In the extended test set, predicted grade was significantly associated with overall survival after adjusting for age and gender (Hazard Ratio 2.05; 95% CI 1.21-3.47); manual grades were not prognostic. Future work can adapt our computational system to predict WHO/ISUP grades, and validating this system on other ccRCC cohorts.

## Introduction

Clear cell renal cell carcinoma (ccRCC) is the most common malignant tumor of epithelial origin in the kidney ^1^. For over 30 years, ccRCC was graded using the 4-tiered Fuhrman nuclear grading system which incorporates nuclear size, nucleolar prominence, and nuclear membrane irregularities. Diagnostic challenges can occur with the presence of other morphological features such as sarcomatoid or spindle cell pattern, when higher grade ccRCC show more eosinophilic staining in the cytoplasm, or in cases where other renal cancer histologic types (e.g. papillary RCC type1 and chromophobe RCC) exhibit clear cytoplasm ^2,3^. The correct classification of ccRCC grade and stage is important to guide clinical management, molecular-based therapies, and prognosis ^4,5^. Fuhrman grade is widely accepted as a prognostic factor despite mediocre inter-observer agreement ^6,7^. To improve inter-observer agreement, simplified 2- or 3-tiered grading systems have been proposed. These simplified systems appear to retain prognostic ability similar to that of 4-tiered systems ^8,9^. Recently, a new nuclear/nucleolar grading system, known as the World Health Organization (WHO)/International Society of Urological Pathology (ISUP) Grading Classification for RCC, was introduced ^10^.

Technological advances have enabled computational pathology to discover novel morphometric features from whole slide images (WSIs) that may add diagnostic and/or prognostic information ^11–13^. Computational pathology techniques can analyze cancer WSIs ^14–16^, including the detection of malignant RCC cells ^17^. In this study, we developed an automated grading system to predict 2-tiered Fuhrman grade using ccRCC WSIs from The Cancer Genome Atlas (TCGA). Our specific aims were to establish a computational pipeline to extract nuclei morphometric features and develop a model to predict 2-tiered ccRCC grade, and evaluate the prognostic efficacy of computer predicted grades.

## Materials and Methods

### Cases and Grade Assignment

TCGA ccRCC clinical data, including Fuhrman’s grade (accessed June 2017), and the hematoxylin and eosin (H&E) WSIs were retrieved for 395 cases ^18,19^. TCGA ccRCC cases were contributed by seven participating medical centers. The TCGA Fuhrman’s grade for each case is the consensus of at least two pathologists from the case’s medical center. In order to identify tumor areas on each diagnostic WSI (i.e., regions of interest (ROIs)) for this computational pathology study, Pathologist 1 reviewed and identified an average of five ROIs for each case. Pathologist 1 assigned a Fuhrman grade of 1 to 4 for each ROI, and the highest grade among all the ROIs was the designated grade. Thus, each patient had two assigned grades: “TCGA grade” and “Grade by Pathologist 1”. TCGA and Pathologist 1 grades were re-stratified into the 2-tiered grading system: low (grades 1 and 2) and high (grades 3 and 4).

### Image Processing, Nuclei Segmentation and Morphometric Feature Extraction

ROIs (*n*=1855) from 395 WSIs were extracted and split into 2000 pixel by 2000 pixel patches. Cell nuclei was segmented using our previously published workflow and Fiji ^20,21^ (Figure 1). Patches were converted from the Red, Green, and Blue (RGB) color space to the Hue, Saturation, and Value (HSV) color space. Adaptive thresholding was performed to identify nuclei regions. Watershed transform separated overlapping nuclei. Extracted nuclei of area less than 200 pixels or greater than 2000 pixels were excluded to improve the specificity of nuclear detection ^14^.

**Figure 1.**
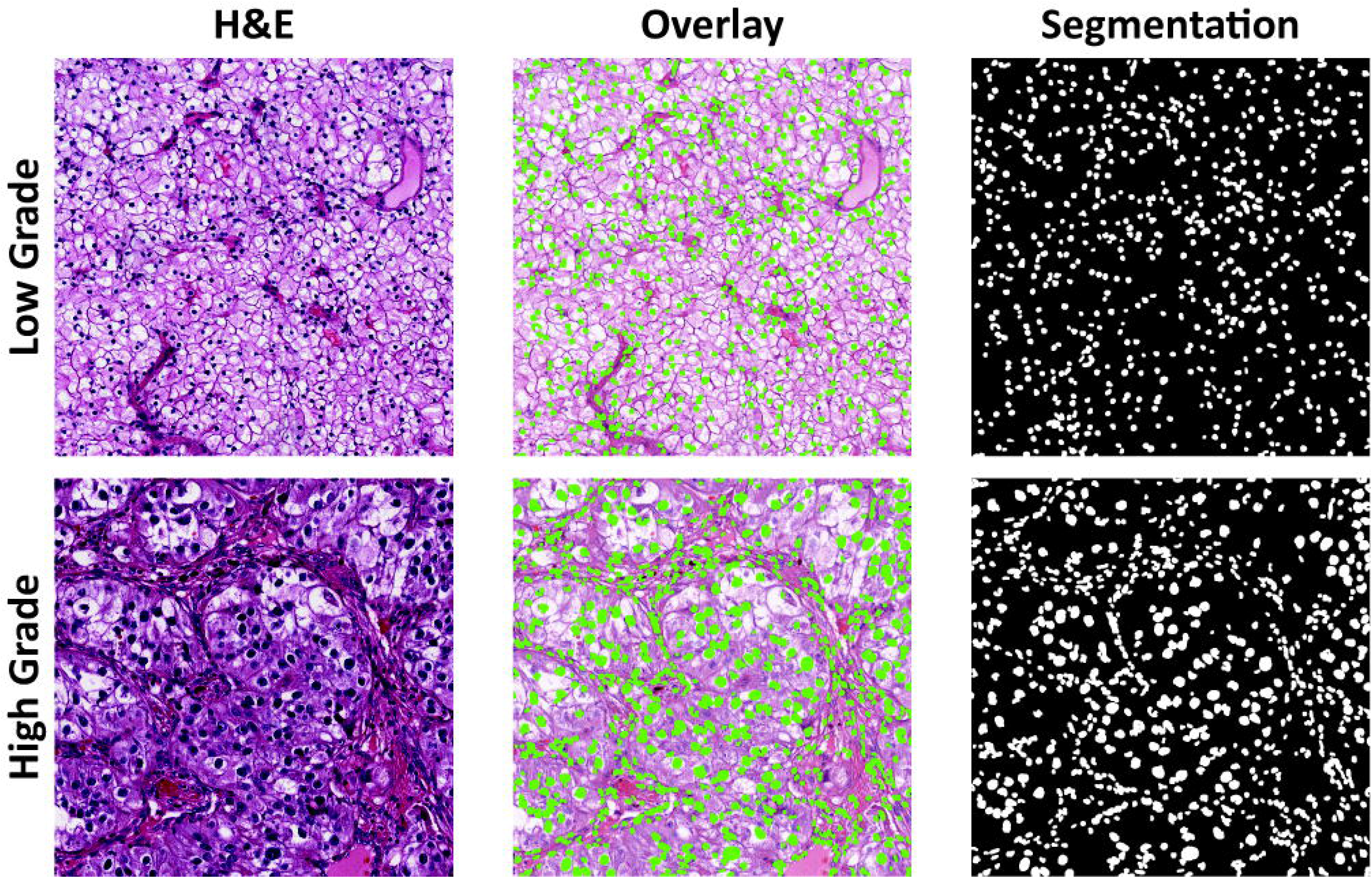
Examples of nuclei detection and segmentation in low and high grade clear cell renal cell carcinoma. The rightmost column shows computer-generated segmentation mask where cell nuclei are labelled white against a black background. The middle column shows the overlay of segmented nuclei (green spots) over each hematoxylin and eosin (H&E) patch.

For each patch, 72 nuclei morphometric features were extracted: nine morphological features, 15 intensity-based features, and 48 texture-based features. Morphological features describe the shape and size variation of nuclei. Intensity features (first order statistical features) describe the distribution of color variation in the nucleus. Three color channels were analyzed: lightness from HSV color space, lightness from Lab color space, and Hematoxylin channel from H&E color deconvolution ^22^. Five first order statistical features were computed—mean, median, standard deviation, skewness, and kurtosis—for each of the three color channels, for a total of 15 intensity features. Texture features (second order statistical features) quantitatively describe patterns and texture of pixel values. Two types of second order statistical features were computed: co-occurrence based features (*n*=8) and run length based features (*n*=8). Co-occurrence based features include correlation, cluster shade, cluster prominence, energy, entropy, Haralick correlation, inertia, and inverse difference moment ^23^. Run length based features include gray-level non-uniformity, run-length non-uniformity, low and high gray-level run emphasis, short run low and high gray-level emphasis, long run low and high gray-level emphasis ^24^. Likewise, texture features were extracted from the three selected color channels, resulting in a total of 48 texture features.

### Data Summarization and Selection of Representative ROI

Data extracted at the patch level were summarized to the ROI level by calculating the median and median absolute deviation (MAD) (i.e., 144 summarized features). Some cases had multiple ROIs annotated with the highest grade. Thus, one ROI among the highest grade ROIs was selected to represent the case. To do so, the median of all ROIs with the highest grade was calculated, and the ROI with the smallest Euclidean-distance to the calculated median was chosen.

### Nuclei Morphometric Features Associated with Grade

Cases with concordant 2-tiered grade by TCGA and Pathologist 1 (*n*=277) were used to develop the automated 2-tiered grading system. Concordant cases were randomly spilt into a training set (*n*=235; 85%) and held-out test set (*n*=42; 15%). Seven machine learning classification methods were explored to classify ccRCC cases into low or high grade using nuclei morphometric features ^25,26^. All methods achieved similar area under the receiver-operator characteristic curves (AUC ROC; Supplementary 1). Lasso regression was the top performing method with a built-in feature selection capability. Furthermore, Lasso is computationally efficient and more interpretable compared to other machine learning methods such as deep learning. Therefore, the final classification model was built using Lasso on the training set using the optimal lambda parameter and evaluated on the test set. This workflow is summarized in Supplementary 2.

### Survival Analyses

The Lasso model was applied to predict the grade of the previously held out test set (*n*=42) and cases with discordant grades (*n*=118). These 160 cases were combined to create an extended test set to evaluate the prognostic capability (i.e., overall survival [OS]) of our predicted grade using crude and adjusted Cox proportional hazard models. The adjusted Cox models include patient age, gender, and cancer stage. TCGA treatment information was missing from 69% of the cases and thus was not included in the adjusted Cox models. Kaplan-Meier curves were plotted to visualize differences between the curves (survival package, R)^27^.

### Additional Pathological Review for Discordant Cases

The grades provided by TCGA may be assessed from ROIs other than the representative ROIs selected in our study. To obtain a fairer comparison between manual and predicted grades among the discordant cases, the representative ROIs were additionally reviewed by Pathologist 2.

### Statistical Analyses

Confusion matrices determined the concordance of the 2-tiered and 4-tiered grades between two raters ^26^. Inter-rater reliability among three raters was evaluated using Fleiss’ kappa. Boxplots were created using ggplot2 version 2.2.1. Comparisons between the nine morphological features with 2-tiered and 4-tiered grading were done using Mann-Whitney U or Kruskal Wallis test, respectively. All tests of statistical significance were two-sided. Statistical significance was achieved when *p*-value was <0.05 or when the false discovery rate (FDR) was <0.05. All analyses were conducted using R version 3.4.0.

## Results

The majority of TCGA ccRCC cases were white males. Most participants were between the ages of 50 to 69 and had stage I disease (Table 1). The agreement of 4-tiered grading between TCGA and Pathologist 1 was poor (frequency of agreement = 0.47, Cohen’s kappa = 0.20; Supplementary 3A). When the grading was stratified into 2-tiers, 277 out of 395 cases were concordant (frequency of agreement = 0.70, Cohen’s kappa = 0.41; Supplementary 3B). Most of the discordant cases were assigned high grade by TCGA and low grade by Pathologist 1.

**Table 1.** Demographic table of the 395 The Cancer Genome Atlas (TCGA) clear cell renal cell carcinoma cases with 2-tiered histological grade (low and high). Note that the TCGA grade for each patient in the discordant set is the opposite grade assigned by Pathologist 1.

|  | Total *n* (%) | Concordant Cases |  | Discordant cases (Grades by TCGA) |  | Discordant cases (Grades by Pathologist 1) |  |
| --- | --- | --- | --- | --- | --- | --- | --- |
|  |  | Low *n* (%) | High *n* (%) | Low *n* (%) | High *n* (%) | Low *n* (%) | High *n* (%) |
| Cases, *n* | 395 (100) | 162 (58.5) | 115 (41.5) | 28 (23.7) | 90 (76.3) | 90 (76.3) | 28 (23.7) |
| Age group, *n* |  |  |  |  |  |  |  |
| <50 | 80 (20.3) | 36 (22.2) | 22 (19.1) | 4 (14.3) | 18 (20.0) | 18 (20.0) | 4 (14.3) |
| 50-59 | 106 (26.8) | 50 (30.9) | 29 (25.2) | 6 (21.4) | 21 (23.3) | 21 (23.3) | 6 (21.4) |
| 60-69 | 109 (27.6) | 37 (22.8) | 33 (28.7) | 10 (35.7) | 29 (32.2) | 29 (32.2) | 10 (35.7) |
| 70-79 | 82 (20.8) | 31 (19.1) | 26 (22.6) | 8 (28.6) | 17 (18.9) | 17 (18.9) | 8 (28.6) |
| >80 | 18 (4.6) | 8 (4.9) | 5 (4.3) | 0 (0.0) | 5 (5.6) | 5 (5.6) | 0 (0.0) |
| Gender, *n* |  |  |  |  |  |  |  |
| Female | 130 (32.9) | 67 (41.4) | 28 (24.3) | 10 (35.7) | 25 (27.8) | 25 (27.8) | 10 (35.7) |
| Male | 265 (67.1) | 95 (58.6) | 87 (75.7) | 18 (64.3) | 65 (72.2) | 65 (72.2) | 18 (64.3) |
| Race, *n* |  |  |  |  |  |  |  |
| Asian | 7 (1.8) | 3 (1.9) | 2 (1.7) | 0 (0.0) | 2 (2.2) | 2 (2.2) | 0 (0.0) |
| Black | 33 (8.4) | 13 (8.0) | 10 (8.7) | 2 (7.1) | 8 (8.9) | 8 (8.9) | 2 (7.1) |
| White | 349 (88.4) | 142 (87.7) | 102 (88.7) | 25 (89.3) | 80 (88.9) | 80 (88.9) | 25 (89.3) |
| Not reported | 6 (1.5) | 4 (2.5) | 1 (0.9) | 1 (3.6) | 0 (0.0) | 0 (0.0) | 1 (3.6) |
| Stage, *n* |  |  |  |  |  |  |  |
| Stage I | 207 (52.4) | 121 (74.7) | 35 (30.4) | 13 (46.4) | 38 (42.2) | 38 (42.2) | 13 (46.4) |
| Stage II | 44 (11.1) | 17 (10.5) | 15 (13.0) | 5 (17.9) | 7 (7.8) | 7 (7.8) | 5 (17.9) |
| Stage III | 92 (23.3) | 18 (11.1) | 36 (31.3) | 8 (28.6) | 30 (33.3) | 30 (33.3) | 8 (28.6) |
| Stage IV | 52 (13.2) | 6 (3.7) | 29 (25.2) | 2 (7.1) | 15 (16.7) | 15 (16.7) | 2 (7.1) |
| Type of Treatment, *n* |  |  |  |  |  |  |  |
| Chemotherapy | 7 (1.8) | 3 (1.9) | 4 (3.5) | 0 (0.0) | 0 (0.0) | 0 (0.0) | 0 (0.0) |
| Immunotherapy | 6 (1.5) | 2 (1.2) | 2 (1.7) | 0 (0.0) | 2 (2.2) | 2 (2.2) | 0 (0.0) |
| Molecular therapy | 79 (20.0) | 31 (19.1) | 24 (20.9) | 5 (17.9) | 19 (21.1) | 19 (21.1) | 5 (17.9) |
| Radiation | 10 (2.5) | 2 (1.2) | 4 (3.5) | 2 (7.1) | 2 (2.2) | 2 (2.2) | 2 (7.1) |
| Mixed therapy | 21 (5.3) | 4 (2.5) | 10 (8.7) | 2 (7.1) | 5 (5.6) | 5 (5.6) | 2 (7.1) |
| Unknown | 272 (68.9) | 120 (74.1) | 71 (61.7) | 19 (67.9) | 62 (68.9) | 62 (68.9) | 19 (67.9) |

Computer extracted morphological features reflected the variation of ccRCC nuclei as observed by pathologists. Nuclei size (i.e., area, perimeter, and spherical perimeter and radius) and shape (i.e., roundness, elongation, flatness and major axis of ellipse fit) were significantly larger and less spherical in higher grades (FDR<0.05; Supplementary 4 and 5).

### Lasso Classification Model

The final Lasso model had an ROC AUC of 0.84. The model predicted 2-tiered ccRCC grade with 84.6% sensitivity, 81.3% specificity, 18.8% false positive rate and 15.4% false negative rate in the test set. The agreement between predicted and manual grades was good (frequency of agreement = 0.83, Cohen’s kappa = 0.65). The 18 unique morphometric features associated with ccRCC 2-tiered grade are in Table 2.

**Table 2.** Nuclear morphometric features associated with 2-tiered ccRCC grade selected in the final Lasso classification model (18 unique features; 26 total features).

| Feature | Type | Color Space | Summary Function | Biological Relevance |
| --- | --- | --- | --- | --- |
| Elongation | Morphology | - | MAD | Nuclear pleomorphism, nuclear shape (irregular) |
| Minor axis of the Ellipse Fit | Morphology | - | Median | Nuclear pleomorphism, nuclear shape (irregular) |
| Flatness | Morphology | - | MAD | Nuclear shape (irregular) |
| Kurtosis | Intensity | HSV | Median | Uneven distribution of nucleus staining |
| Skewness | Intensity | H&E | MAD | Uneven distribution of nucleus staining |
|  |  | HSV | Median |  |
|  |  | Lab | MAD |  |
| Correlation | Texture | HSV | MAD | Granularity of chromatin (a) |
| Haralick Correlation | Texture | H&E | Median | Granularity of chromatin (a) |
| Energy | Texture | Lab | Median & MAD | Granularity of chromatin (b) |
| Inverse difference moment | Texture | H&E | Median & MAD | Granularity of chromatin (b) |
|  |  | HSV | Median & MAD |  |
| Inertia | Texture | H&E | Median | Granularity of chromatin (b) |
|  |  | Lab | Median |  |
| Entropy | Texture | HSV | Median | Granularity of chromatin (c) |
| Low gray-level run emphasis | Texture | H&E | MAD | Granularity of chromatin (c) |
| Long run high gray-level emphasis | Texture | HSV | MAD | Granularity of chromatin (c) |
| Long run low gray-level emphasis | Texture | H&E | MAD | Granularity of chromatin (c) |
| Short run high gray-level emphasis | Texture | HSV | MAD | Granularity of chromatin (c) |
| Short run low gray-level emphasis | Texture | H&E | MAD | Granularity of chromatin (c) |
| Gray level non-uniformity | Texture | HSV | Median | Granularity of chromatin (d) |
|  |  | Lab | MAD |  |
| High gray-level run emphasis | Texture | HSV | MAD | Granularity of chromatin (d) |

- a) Correlation is a co-occurrence based texture feature, describing roughness and repeated direction inside the nuclei. - b) Co-occurrence based texture feature, describing roughness inside the nuclei. - c) Co-occurrence based texture feature, describing randomness of gray-level distribution. - d) Run-length based texture feature, describing coarseness inside nuclei.
MAD: median absolute deviation; Lab: Lab color space; HSV: hue-saturation-value color space; H&E: Hematoxylin and Eosin color space

### Prognostic Efficacy of Predicted Grades

There were 65 death events out of 160 cases in the extended test set. Cases predicted as high grade had significantly poorer OS compared to low grade (Figure 2). The association between predicted grade and OS was significant in the crude analysis (hazard ratio (HR) 2.07; 95% confidence interval (CI) 1.25-3.43) and after adjusting for age and gender (HR 2.05; 95% CI 1.21-3.47). The association was attenuated when stage was included in the model (HR 1.66; 95% CI 0.97-2.83).

**Figure 2.**
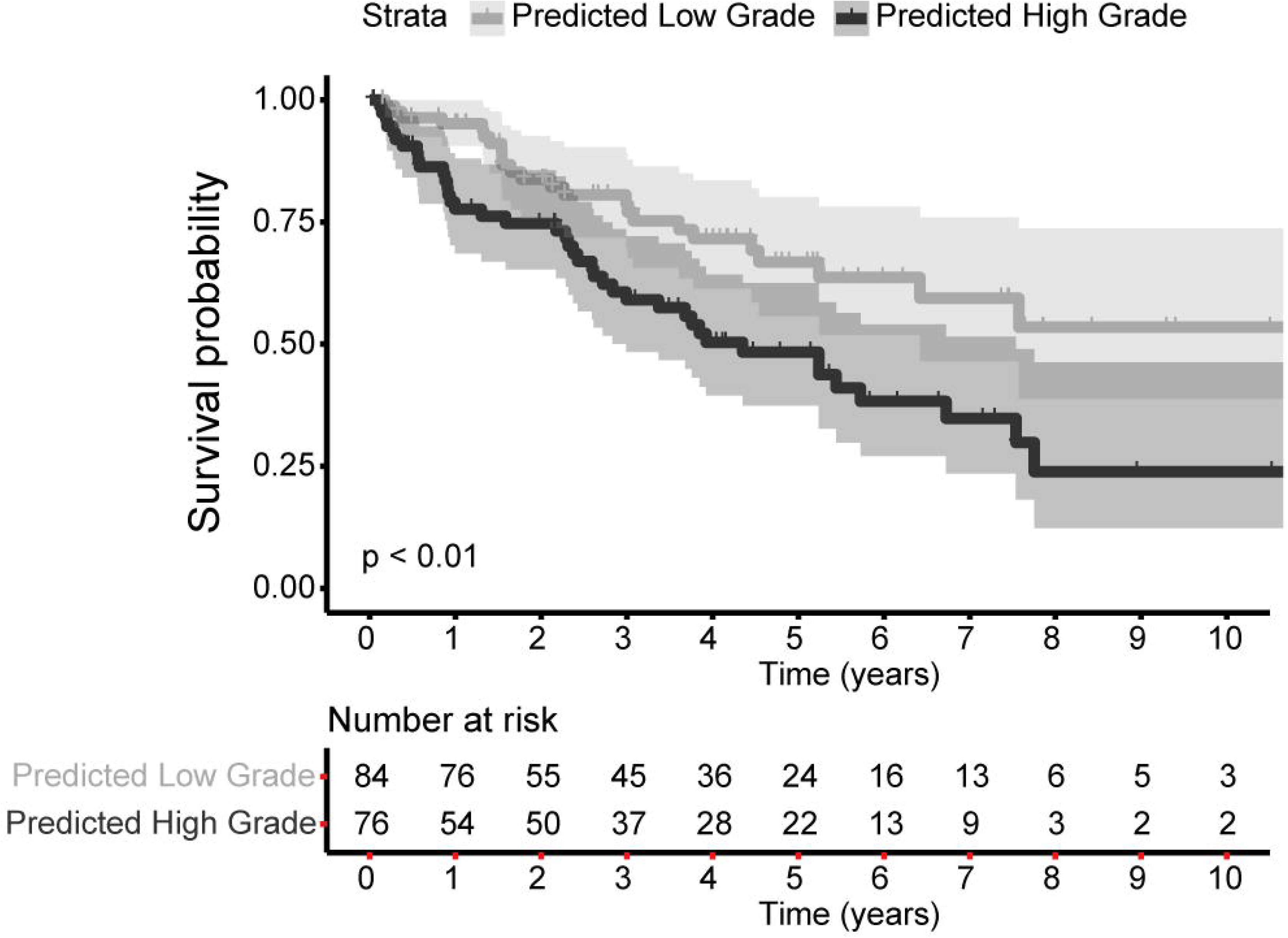
Cases predicted as high grade have significantly poorer overall survival rates compared to cases predicted as low grade in the extended test set (hazard ratio 2.07, 95% confidence interval of 1.25-3.43, *p*<0.01; 65 death events among 160 cases). The shaded areas reflect the 95% confidence interval for high or low grade.

### Comparing Predicted Grade with TCGA and Pathologist 1

Among the concordant cases, 2-tiered manual grades were significantly associated with OS (Figure 3A; Table 3). Predicted grade for concordant cases were not evaluated as the majority of the concordant cases were part of the training set used to build the Lasso model. Within the discordant cases, neither grade provided by TCGA nor Pathologist 1 was associated with OS (Figures 3B and 3C). Predicted grade was significantly associated with OS (crude model HR 2.01; 95% CI 1.14-3.54) and when adjusted for age and gender (HR 2.31; 95% CI 1.26-4.24). The association of predicted grade and OS among the discordant cases was attenuated when adjusted stage was included in the model (HR 1.83; 95% CI 0.98-3.41; Figure 3D; Table 3).

**Figure 3.**
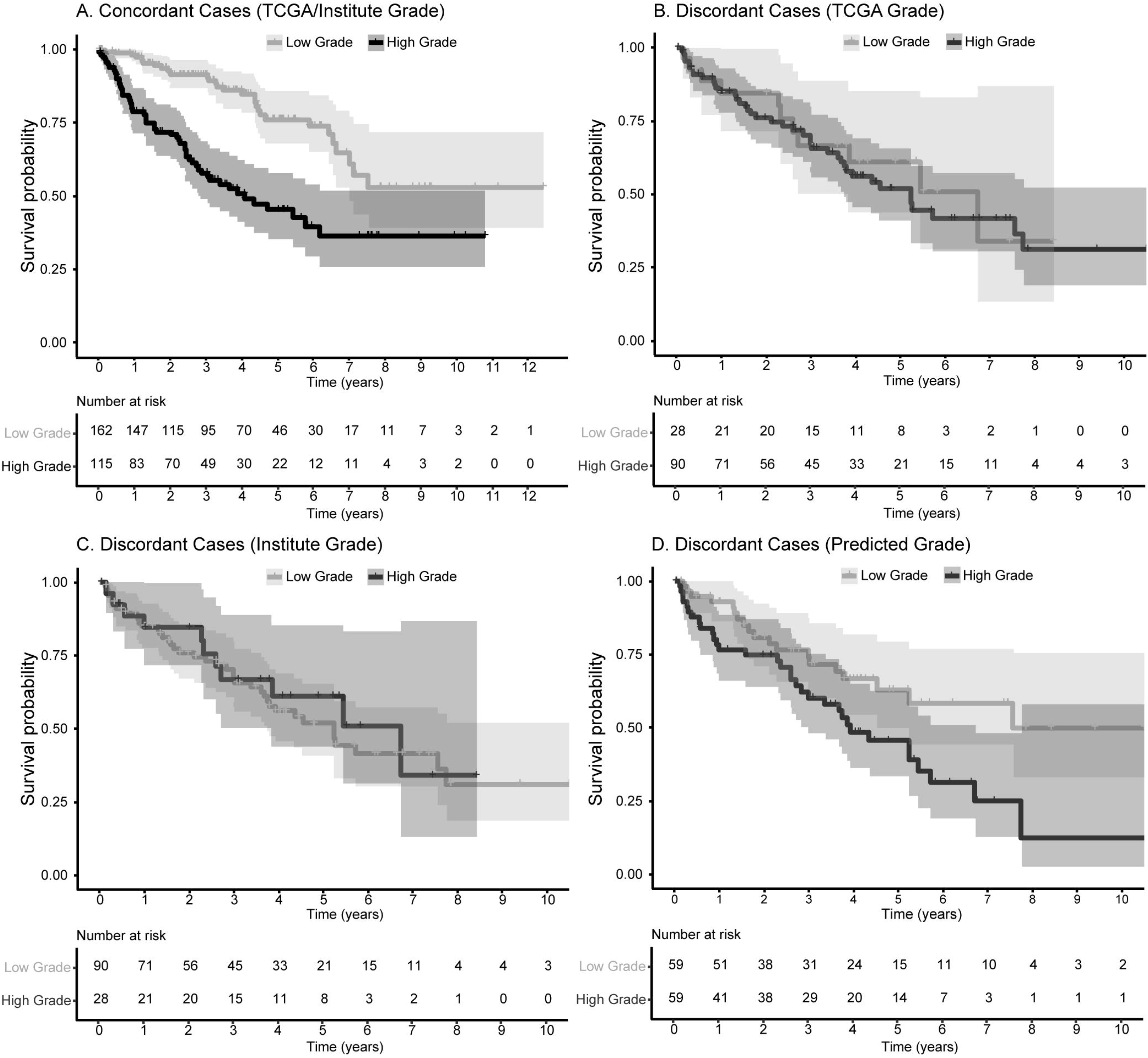
This set of Kaplan-Meier curves compared manual and predicted grades with overall survival in concordant and discordant cases. Grades assigned by TCGA/Pathologist 1 were significantly associated with overall survival within the concordant cases (**A**). In the discordant set, neither grades assigned by TCGA (**B**) nor Pathologist 1 (**C**) were associated with overall survival while predicted grade remained significantly prognostic (**D**). Please refer to Table 3 for hazard ratios and 95% confidence intervals for each analysis. The shaded areas reflect the 95% confidence interval for high or low grade.

**Table 3.** The association of manual or computer predicted 2-tiered grade with overall survival in the concordant and discordant cases.

|  | Manual Grade |  |  | Computer Predicted Grade |  |  |
| --- | --- | --- | --- | --- | --- | --- |
|  | Hazard Ratio | (95% CI) | p-value | Hazard Ratio | (95% CI) | p-value |
| **A. Concordant cases between TCGA and Pathologist 1 (85 events out of 277 cases)** |  |  |  |  |  |  |
| Model A: Crude | 3.12 | (2.00, 4.86) | **<0.01** | NA | NA | NA |
| Model B: Adjusted for Age and Gender | 3.00 | (1.91, 4.71) | **<0.01** | NA | NA | NA |
| Model C: Adjusted for Age, Gender, and Stage | 1.59 | (0.99, 2.57) | 0.06 | NA | NA | NA |
| **B. Discordant cases between TCGA and Pathologist 1 (52 events out of 118 cases)** |  |  |  |  |  |  |
| **Grade assigned by TCGA** |  |  |  |  |  |  |
| Model A: Crude | 1.16 | (0.59, 2.25) | 0.67 | 2.01 | (1.14, 3.54) | **0.02** |
| Model B: Adjusted for Age and Gender | 1.09 | (0.56, 2.13) | 0.80 | 2.31 | (1.26, 4.24) | **<0.01** |
| Model C: Adjusted for Age, Gender, and Stage | 1.08 | (0.55, 2.11) | 0.83 | 1.83 | (0.98, 3.41) | 0.06 |
| **Grade assigned by Pathologist 1** |  |  |  |  |  |  |
| Model A: Crude | 0.86 | (0.44, 1.68) | 0.67 | NA | NA | NA |
| Model B: Adjusted for Age and Gender | 0.92 | (0.47, 1.79) | 0.80 | NA | NA | NA |
| Model C: Adjusted for Age, Gender, and Stage | 0.93 | (0.47, 1.82) | 0.83 | NA | NA | NA |
| **Grade assigned by Pathologist 2** |  |  |  |  |  |  |
| Model A: Crude | 1.09 | (0.63, 1.89) | 0.75 | NA | NA | NA |
| Model B: Adjusted for Age and Gender | 1.21 | (0.70, 2.10) | 0.50 | NA | NA | NA |
| Model C: Adjusted for Age, Gender, and Stage | 1.15 | (0.66, 2.00) | 0.62 | NA | NA | NA |

Confidence Interval, CI

### Additional Pathological Review for Discordant Cases

There was no effective agreement between TCGA, Pathologist 1 and Pathologist 2 among the discordant cases (4-tiered grading: Fleiss’ kappa = −0.23; 2-tiered grading: Fleiss’ kappa = −0.33). When comparing between TCGA and Pathologist 2, there was no effective agreement (4-tiered grading: frequency of agreement = 0.33, Cohen’s kappa = −0.14; 2-tiered grading: frequency of agreement = 0.39, Cohen’s kappa = −0.19). Despite assessing the same representative ROIs, the agreement between Pathologist 1 and Pathologist 2 was poor for 4-tiered grading (frequency of agreement = 0.48, Cohen’s kappa = 0.11) and was slightly improved for 2-tiered grading (frequency of agreement=0.61, Cohen’s kappa = 0.20). Discordant cases between Pathologist 1 and Pathologist 2 were more likely to be assigned as high grade by Pathologist 2. Contingency tables between TCGA, Pathologist 1, and Pathologist 2 are in Supplementary 6.

Grades assigned by Pathologist 2 were not associated with OS (Table 3). Further analyses were explored to determine if the incorporation of manual grade by Pathologist 2 may improve prognostic efficacy. The grades for discordant cases were re-assigned as low or high by using the most frequent grade among TCGA, Pathologist 1, and Pathologist 2, and among Pathologist 1, Pathologist 2, and predicted grade (i.e., integrating manual and computer). Re-assigned grades were not associated with OS (*p*>0.05; Supplementary 7). Next, these cases were spilt into cases that did and did not agree between Pathologist 1 and Pathologist 2. Manual grades were not associated with OS in cases that did and did not agree between Pathologist 1 and Pathologist 2 (*p*>0.05; Table 4). Predicted grade was only associated with OS in cases that agreed between Pathologist 1 and Pathologist 2 (Table 4). Supplementary 8 contains the manual and predicted grades of these ccRCC cases.

**Table 4.** The association of manual or computer predicted 2-tiered grade with overall survival in 118 discordant cases.

|  | Manual Grade |  |  | Computer Predicted Grade |  |  |
| --- | --- | --- | --- | --- | --- | --- |
|  | Hazard Ratio | (95% CI) | p-value | Hazard Ratio | (95% CI) | p-value |
| **A. Cases with identical grades between Pathologist 1 and Pathologist 2 (31 events out of 76 cases)** |  |  |  |  |  |  |
| Model A: Crude | 0.99 | (0.44, 2.23) | 0.99 | 2.05 | (1.00, 4.21) | **0.05** |
| Model B: Adjusted for Age and Gender | 1.04 | (0.46, 2.34) | 0.93 | 2.42 | (1.13, 5.20) | **0.02** |
| Model C: Adjusted for Age, Gender, and Stage | 1.18 | (0.52, 2.68) | 0.69 | 1.89 | (0.87, 4.12) | 0.11 |
| **B. Cases with different grades between Pathologist 1 and Pathologist 2 (21 events out of 46 cases)** |  |  |  |  |  |  |
| **Grade assigned by Pathologist 1** |  |  |  |  |  |  |
| Model A: Crude | 0.64 | (0.19, 2.20) | 0.48 | 2.49 | (0.83, 7.45) | 0.10 |
| Model B: Adjusted for Age and Gender | 0.63 | (0.18, 2.17) | 0.46 | 2.49 | (0.72, 7.28) | 0.16 |
| Model C: Adjusted for Age, Gender, and Stage | 0.59 | (0.17, 2.03) | 0.40 | 2.03 | (0.62, 6.66) | 0.24 |
| **Grade assigned by Pathologist 2** |  |  |  |  |  |  |
| Model A: Crude | 1.56 | (0.46, 5.31) | 0.48 | NA | NA | NA |
| Model B: Adjusted for Age and Gender | 1.58 | (0.46, 5.44) | 0.46 | NA | NA | NA |
| Model C: Adjusted for Age, Gender, and Stage | 1.70 | (0.49, 5.89) | 0.40 | NA | NA | NA |

Confidence Interval, CI

## Discussion

This study utilized the large and diverse TCGA ccRCC dataset to extract quantitative morphometric features from ROIs and applied machine learning algorithms to develop an automated 2-tiered grading system. Using discordant cases as an independent validation set, our data-driven system stratified ccRCC cases into low and high grades that were significantly associated with OS; the prognostic efficacy of predicted grades outperformed the manual grades assessed by TCGA, Pathologist 1, and Pathologist 2. This proof-of-concept study demonstrated the potential of computational pathology to predict ccRCC grades via a more objective and quantitative pipeline, as well as address the issue of grade disagreement commonly encountered between pathologists.

The grading of ccRCC is highly challenging and subjective, but the accurate assignment of ccRCC grade is important for clinical care and follow-up. To the best of our knowledge, this is the first computational pathology system created to predict 2-tiered ccRCC grade with prognostic significance. Previously, Yeh and colleagues developed an automated system to predict grade using a dataset of 39 patients and one feature (i.e., maximum nuclei size), and correlated their predicted grade with manual grade assessed by one pathologist ^28^. Our system, trained using a much larger and more diverse dataset of 277 cases from seven TCGA participating institutions, captures 72 nuclei details in addition to morphological features (i.e., nuclei size and shape) typically observed by pathologists. We showed that computer extracted morphological features were significantly associated with grade. We identified 18 unique morphometric biomarkers to accurately classify ccRCC. These 18 features collectively describe the nucleus, the uneven distribution of nucleus staining, and the granularity of chromatin and nucleoli. This highlights that the addition of computer textual and intensity-related features to morphological features can improve the ability to predict ccRCC grade that correspond to Fuhrman’s grading system and with prognostic utility.

Each TCGA grade is the consensus of at least two pathologists. One reason for grade disagreement between TCGA and Pathologist 1 can be explained by TCGA pathologists assessing different ROIs from the representative ROIs selected in our study. However, even when reviewing the same ROIs for discordant cases, there was very poor agreement between Pathologist 1 and Pathologist 2, reiterating the challenges of ccRCC grading. These discordant cases could be more diagnostically challenging or ambiguous. Since manual grades for concordant cases were significantly associated with OS, it could be argued that concordant cases were diagnostically less challenging where the tumors were overwhelmingly of a low or high grade and that our model was trained using more homogeneous ROIs. Predicted grades for discordant cases were significantly associated with survival, in contrast to manual assessments or using the most frequent manual grade. Therefore, our automated system has the ability to diagnose a range of ccRCC cases with consistency and objectivity. In practical application, such computational system could be useful as a tool to provide a second-opinion in diagnostically ambiguous cases for pathologists.

Our study has some limitations. We did not use the WHO/ISUP grading system because the TCGA participating medical centers used the Fuhrman’s system. However, since our computer system was constructed based on computer extracted nuclear features, it can be adapted to predict WHO/ISUP grades which also utilize nuclei/nucleoli features in the future. There are inherent limitations of reviewing cases using WSIs. Accurate grading may be hindered by the quality of WSIs and the lack of the Z-axis ^29^. Our study reviewed diagnostic WSIs and analyzed manually selected ROIs that may not be representative of the entire tumor. For future work, automating ROI detection and grade prediction will allow the review of multiple tumor sections more efficiently. Lastly, our nuclei segmentation relied on conventional image analysis techniques. While qualitative evaluation of the segmentation results revealed that our image processing pipeline produced reasonably good results, the nuclei segmentation may not be optimal in more challenging cases. A solution is to employ deep learning based techniques to improve nuclei segmentation in future studies ^28,30,31^.

## Conclusions

We developed an automated 2-tiered Fuhrman’s grading system with prognostic significance. Our system demonstrated the potential of computational pathology to improve the reproducibility in the diagnosis and grading of ccRCC, and aid the clinical management of ccRCC patients. Future work may include adapting our computational system to predict WHO/ISUP grades; validating our system on other ccRCC cohorts; using deep learning methods to detect ROIs, segment nuclei and predict grade; and exploring whether morphometric features can predict prognosis independently of grade. This work is one step toward developing an artificial intelligence system for diagnostic pathology.

## Authors’ contributions

Conceived and designed the study: HI, YJH and KT. Image processing and feature extraction: HI and MEP. Data analyses: KT, MV and YJH. Case review: CAR and DIL. All authors contributed to the writing and reviewing of the manuscript.

## Acknowledgements

The data used in this study were in whole or in part based on the data generated by the TCGA Research Network: http://cancergenome.nih.gov/. YJH is supported by the Klarman Family Foundation. KT was supported by the BIDMC High School Summer Research Program.

## Data Availability Statement

TCGA data are available at http://cancergenome.nih.gov/. Supplementary 8 contains the manual and predicted grades of these participants.

