## Supplementary Materials 1-7 for "Automated Clear Cell Renal Carcinoma Grade Classification with Prognostic Significance"

**Supplementary 1.** The average area under the receiver-operator characteristic curves (AUC ROC) for each machine learning method using the training set after 100 iterations of random 10% hold out. These methods were implemented using the glmnet and caret packages in R.

| Rank | Method | AUC ROC using the training set |
| --- | --- | --- |
| 1 | Ensemble | 0.839 |
| 2 | Lasso | 0.834 |
| 3 | Elastic net | 0.823 |
| 4 | Ridge | 0.809 |
| 5 | Neural network | 0.800 |
| 6 | Linear support vector machine | 0.799 |
| 7 | Random forest | 0.781 |

**Supplementary 2.** A summary of the workflow used to develop the 2-tiered clear cell renal cell carcinoma (ccRCC) grade classification. Seven machine learning classification methods were evaluated to determine the optimal method to develop a robust classification model for ccRCC using cases from the Training Set (**A**). Lasso regression produced an average area under the receiver operator characteristic curve of 0.84 and identified nuclei morphometric features associated with ccRCC grade. The Test Set was used to evaluate the performance of the final model; and grades were predicted in the Extended Test Set (**B**).

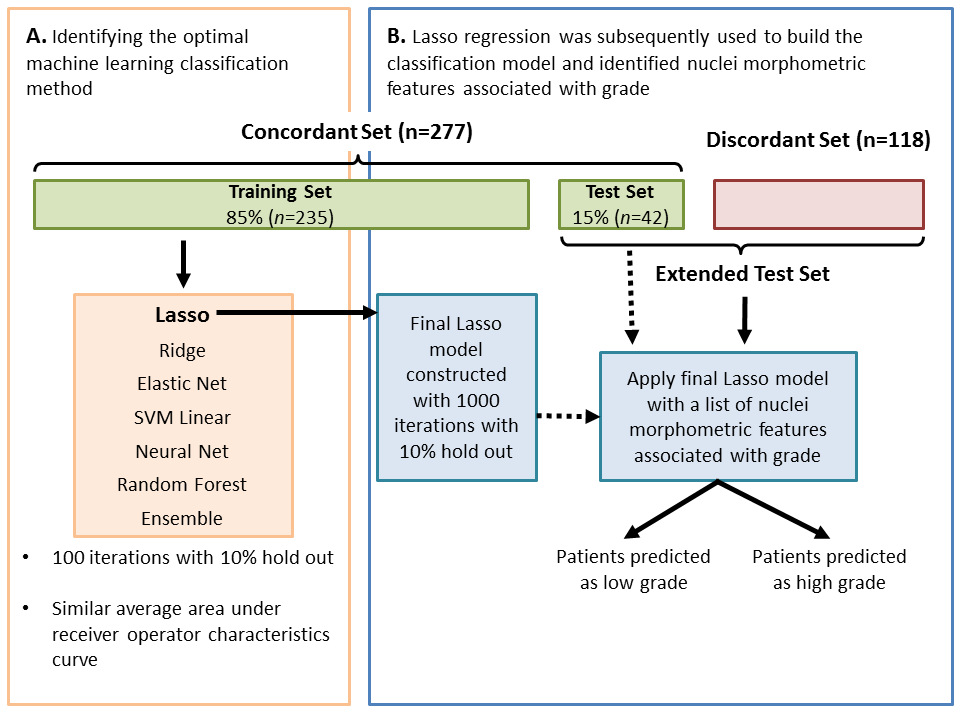

**Supplementary 3. (A)** The agreement between the TCGA and Pathologist 1 using the 4-tiered grading was poor (frequency of agreement = 0.47, Cohen’s kappa = 0.20). **(B)** The agreement improved to moderate when using the 2-tiered grading system (frequency of agreement = 0.70, Cohen’s kappa = 0.41).

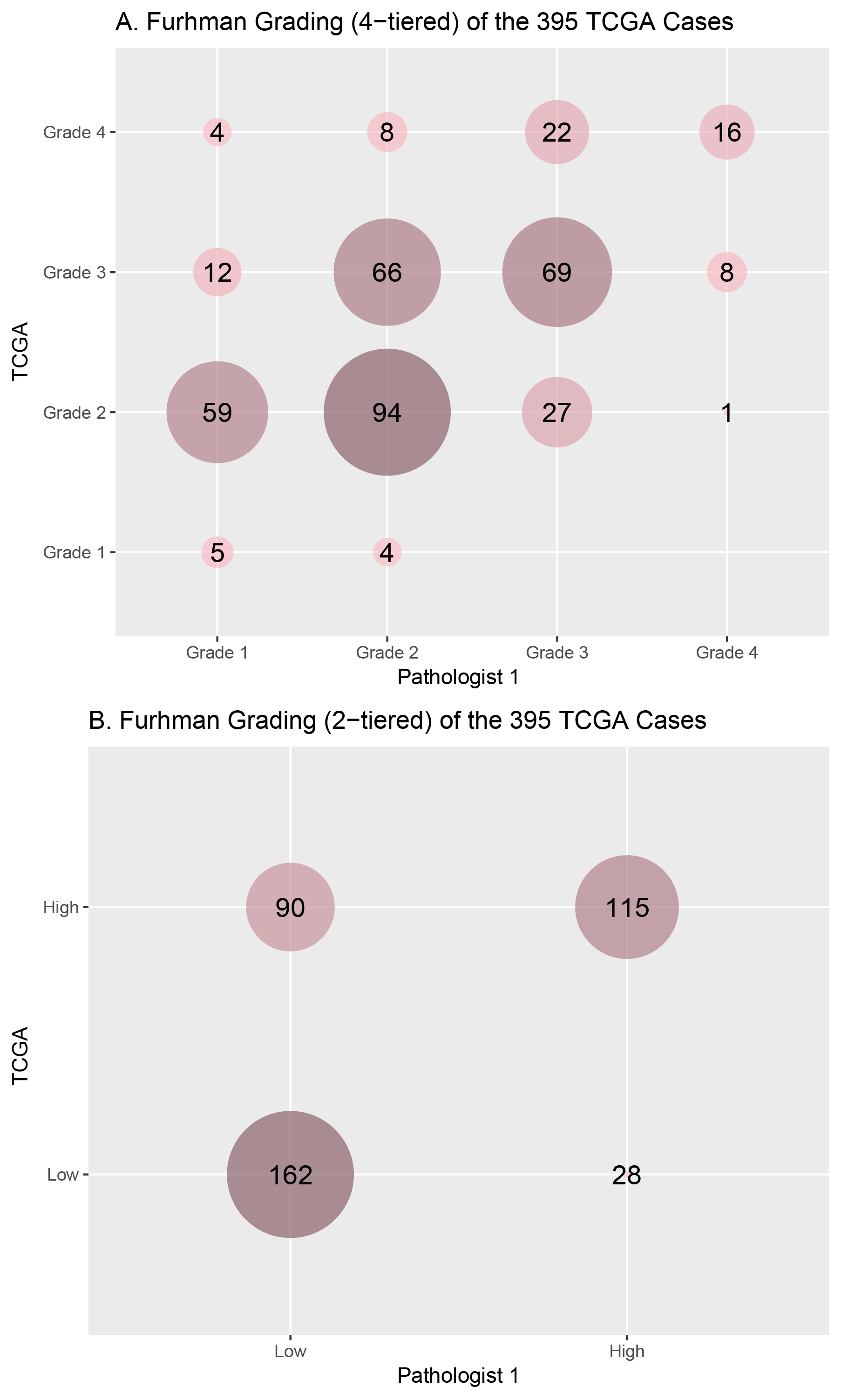

**Supplementary 4.** These plots contain the 277 cases that were concordant between TCGA and Pathologist 1 using the 2-tiered grading system. Nine morphological features (**1:** Area, 2: Roundness, **3:** Elongation, **4:** Flatness, **5:** Perimeter, **6:** Equivalent Spherical Perimeter, **7:** Equivalent Spherical Radius, **8:** Minor Axis of the Ellipse Fit, and **9:** Nuclei Major Axis of the Ellipse Fit) were plotted with their median and median absolute deviation values. Each feature and measure was stratified either by the 4-tiered (**A and B**) or 2-tiered (**C and D**) grades assigned by Pathologist 1, respectively.
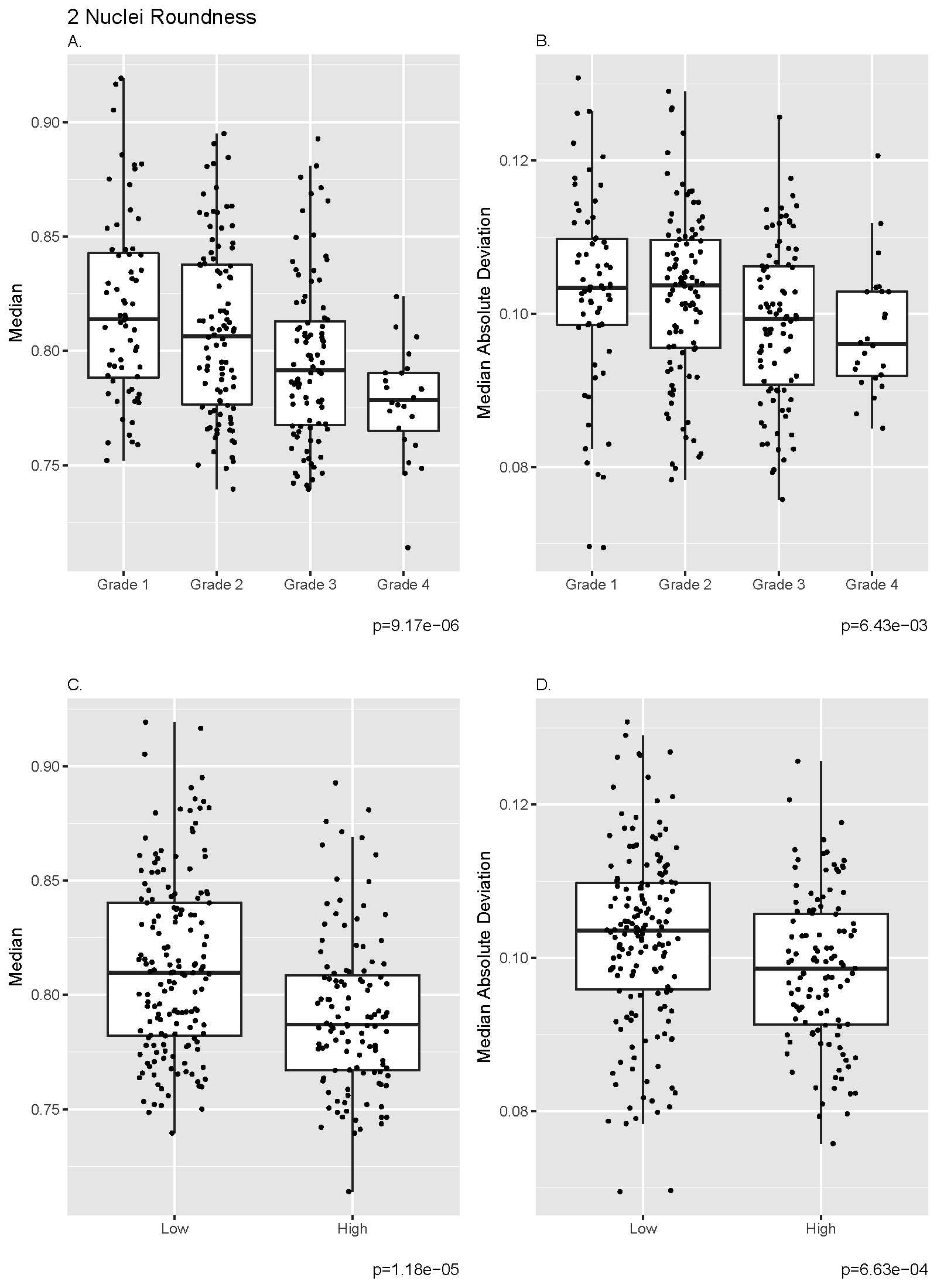

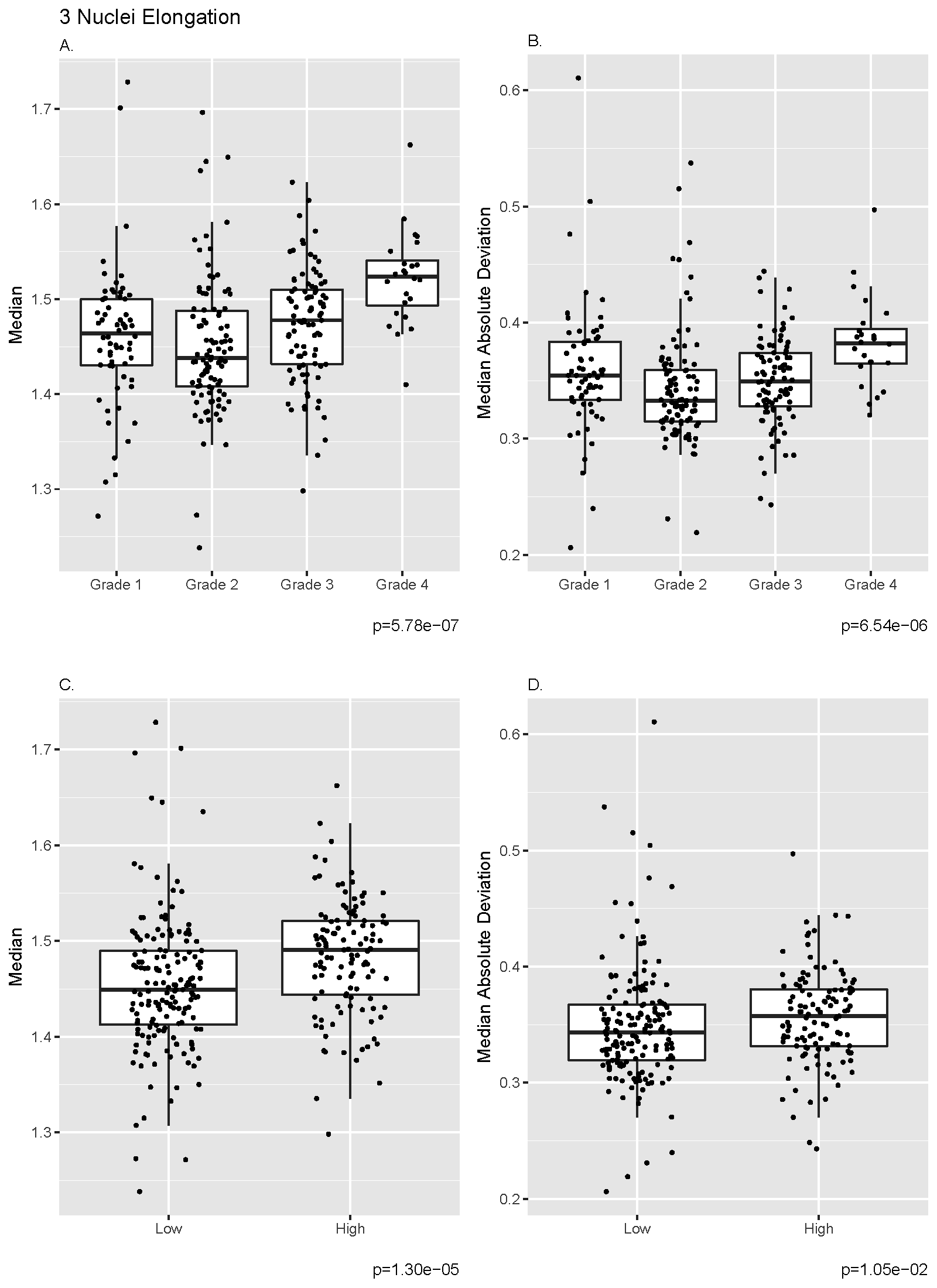

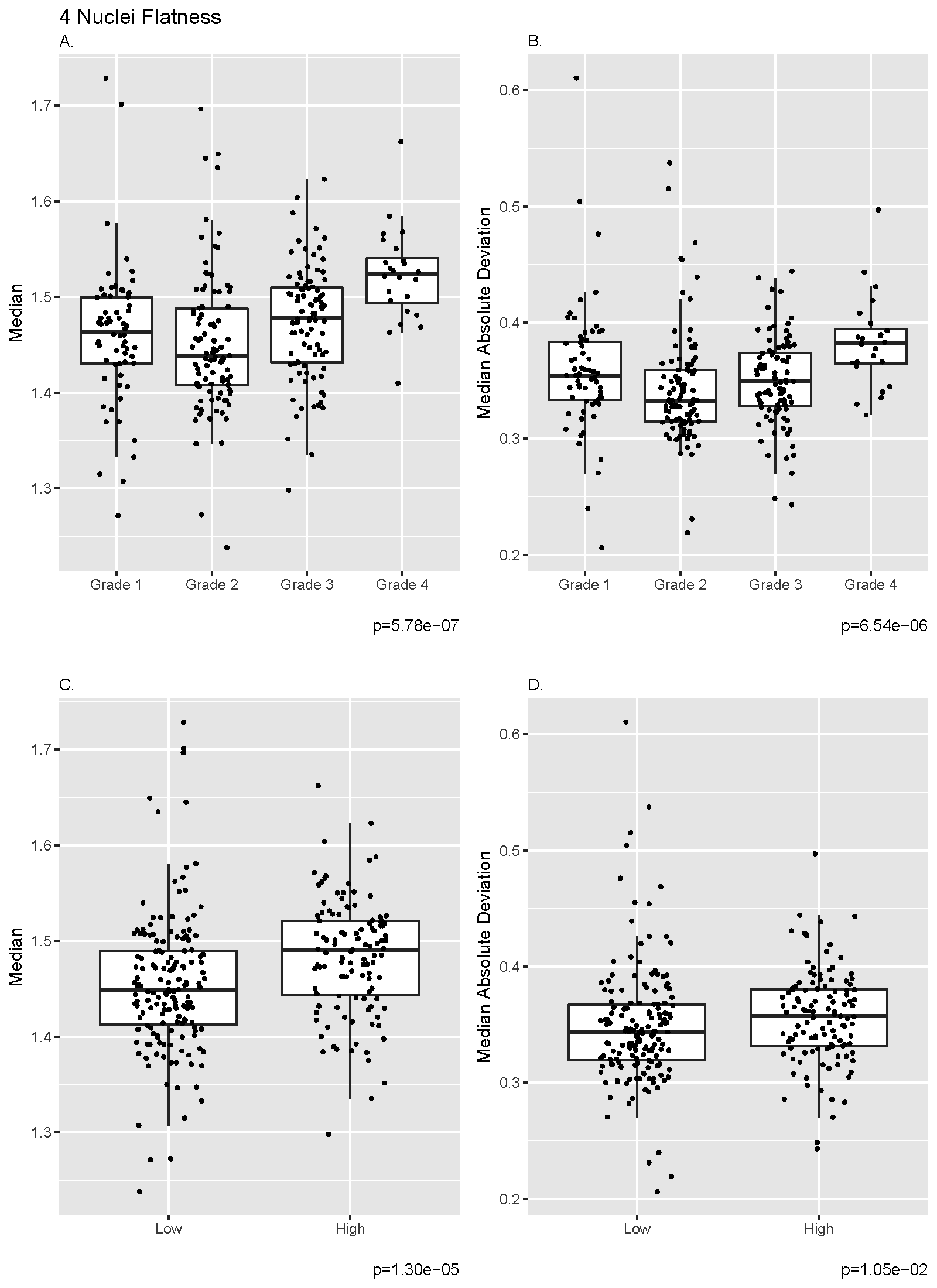

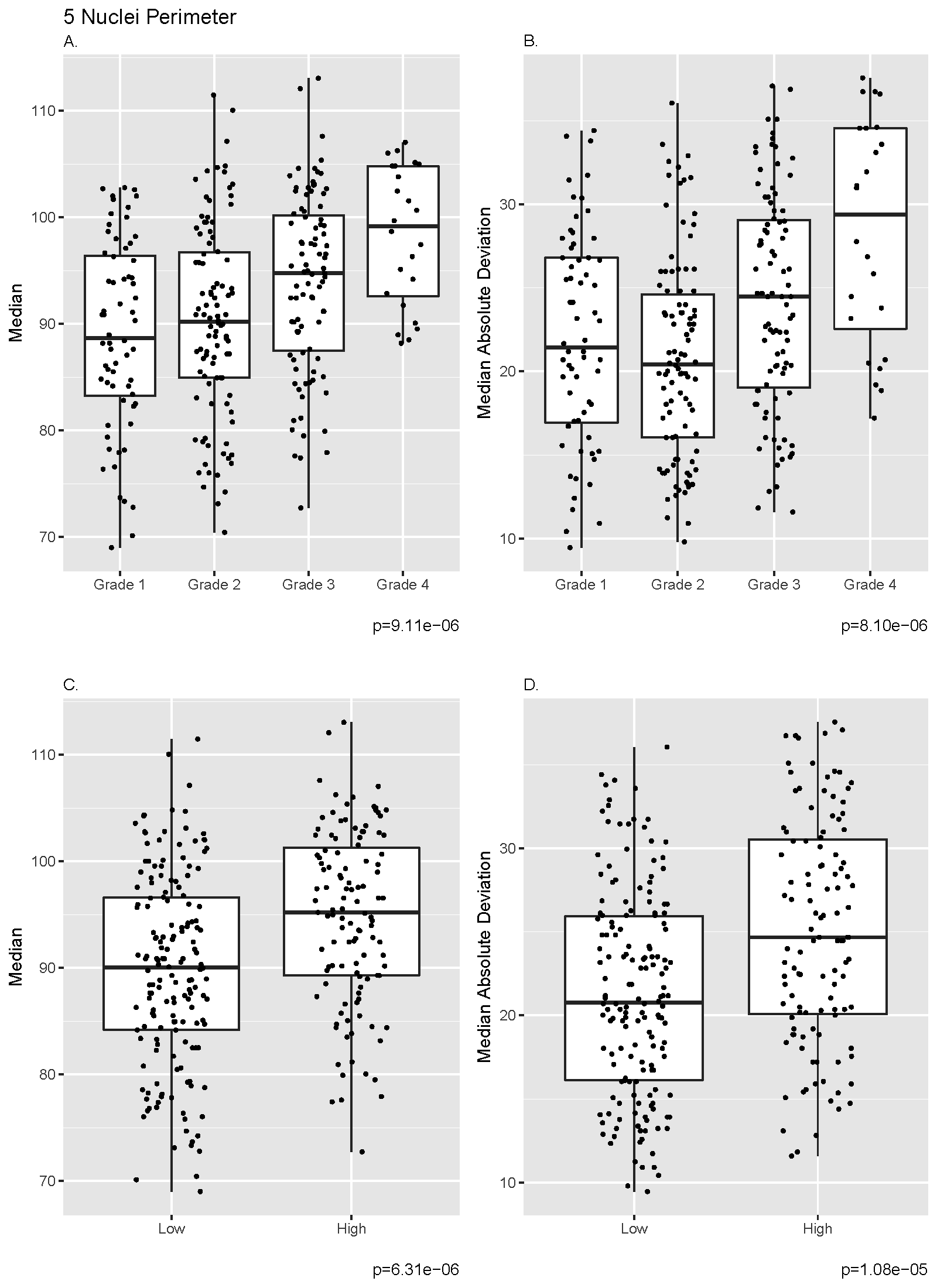

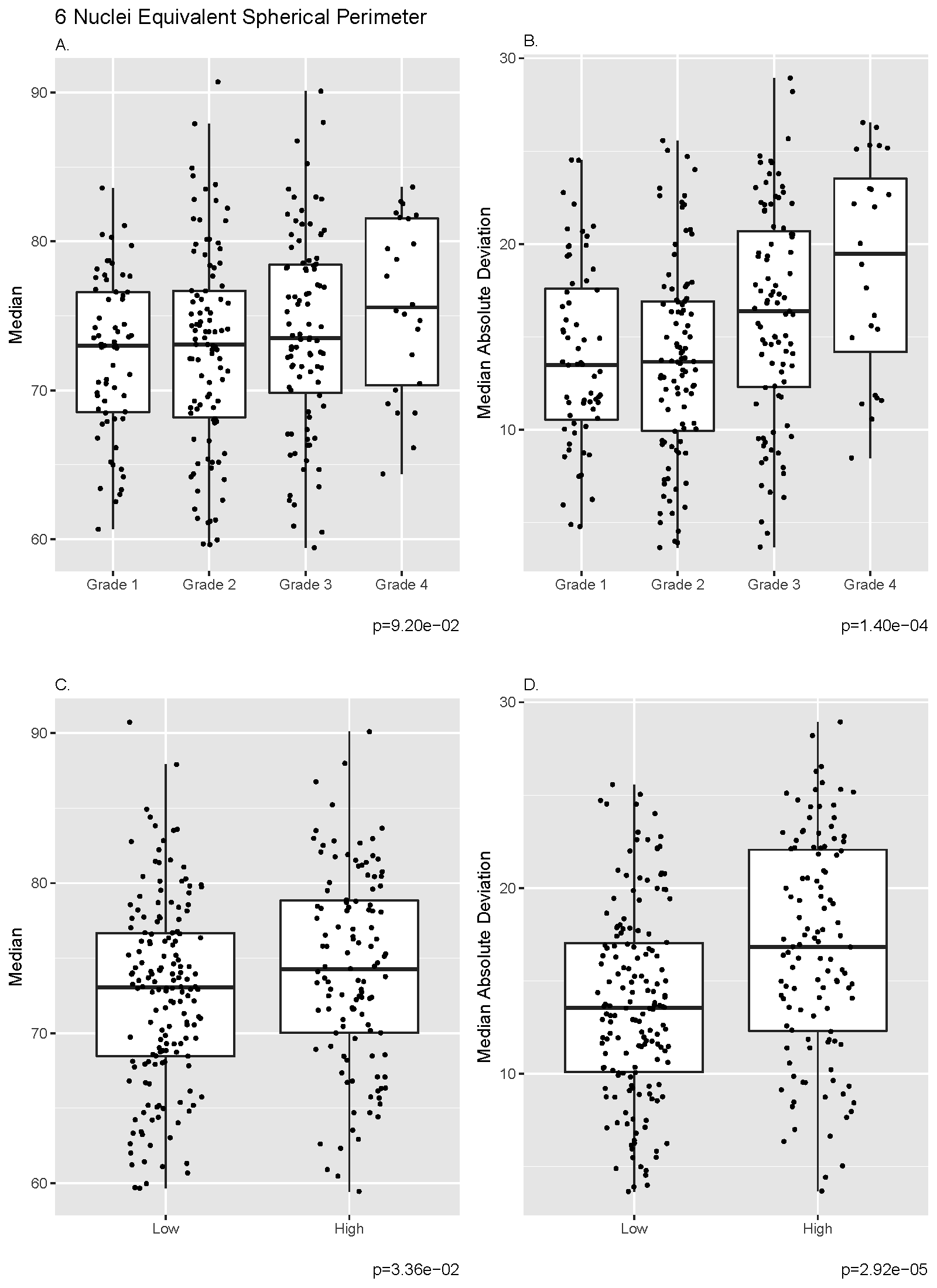

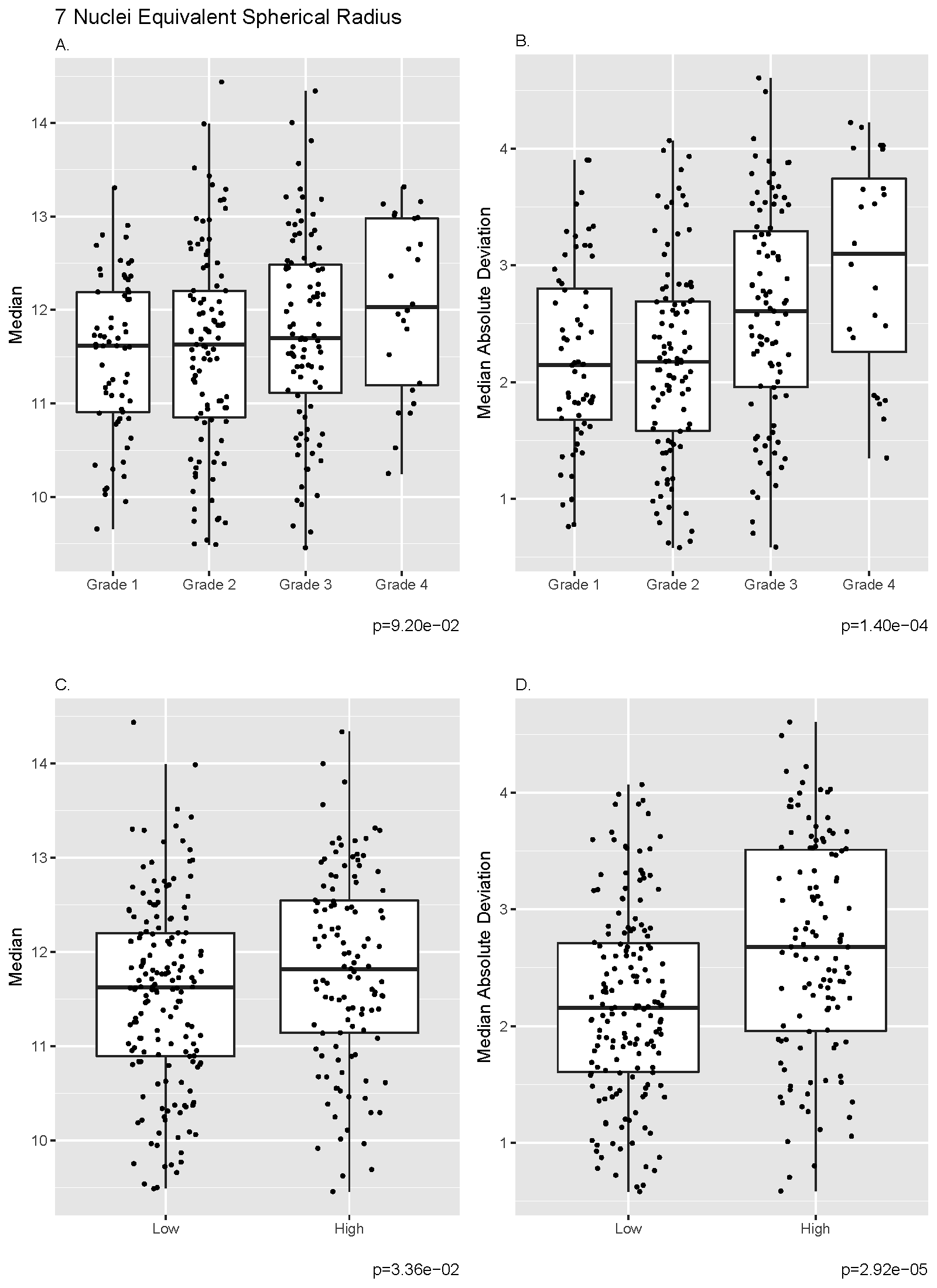

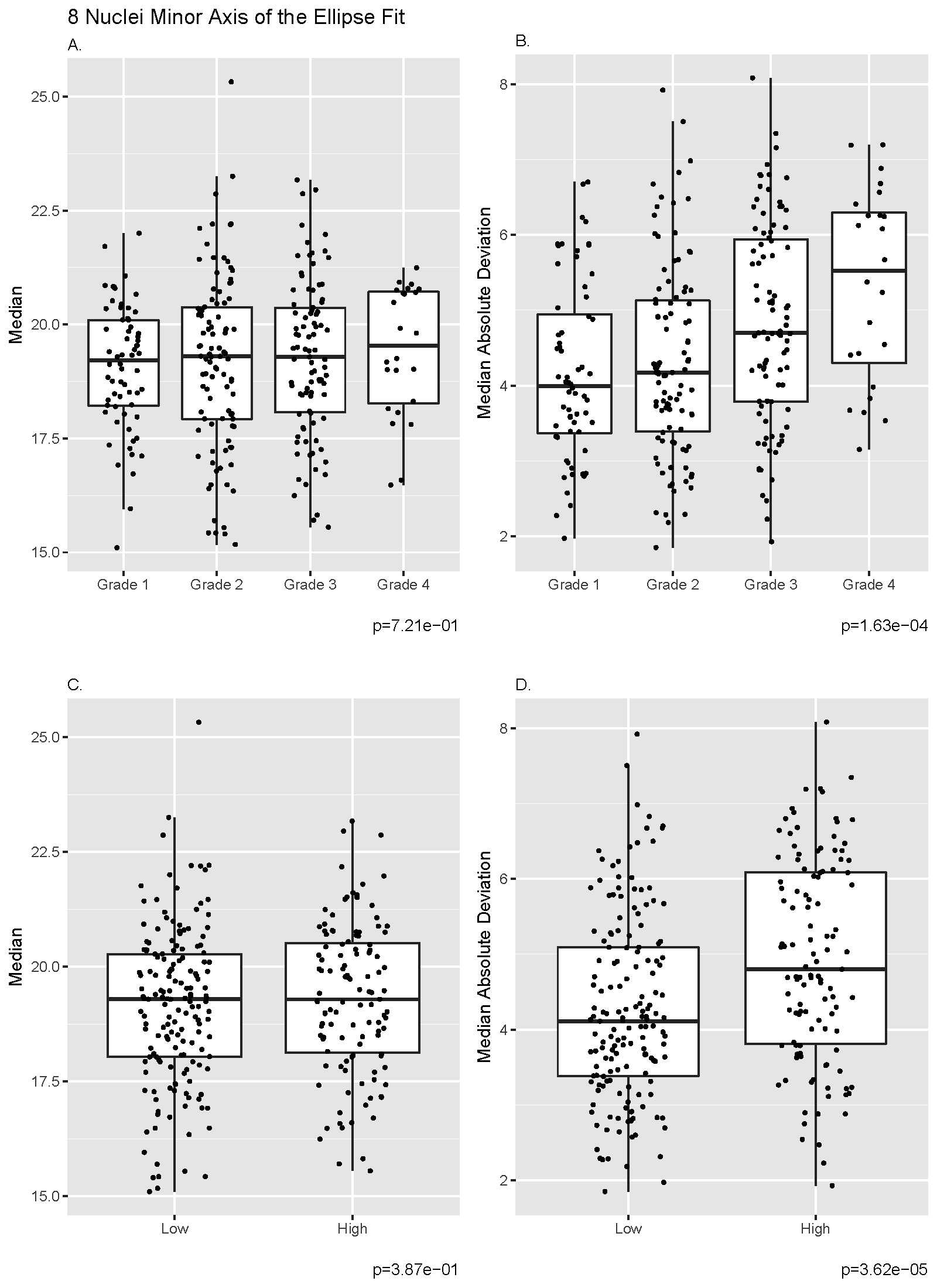

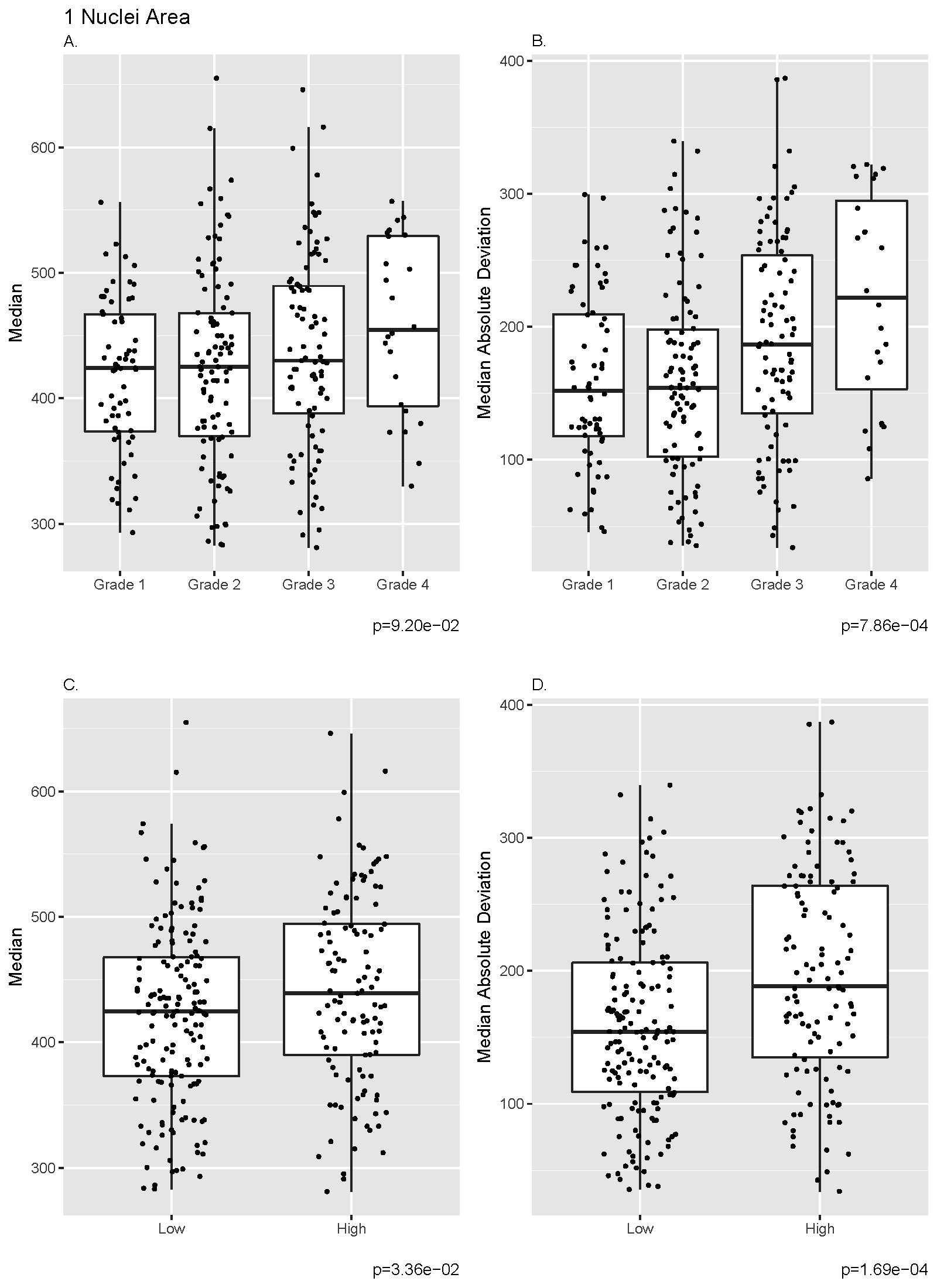

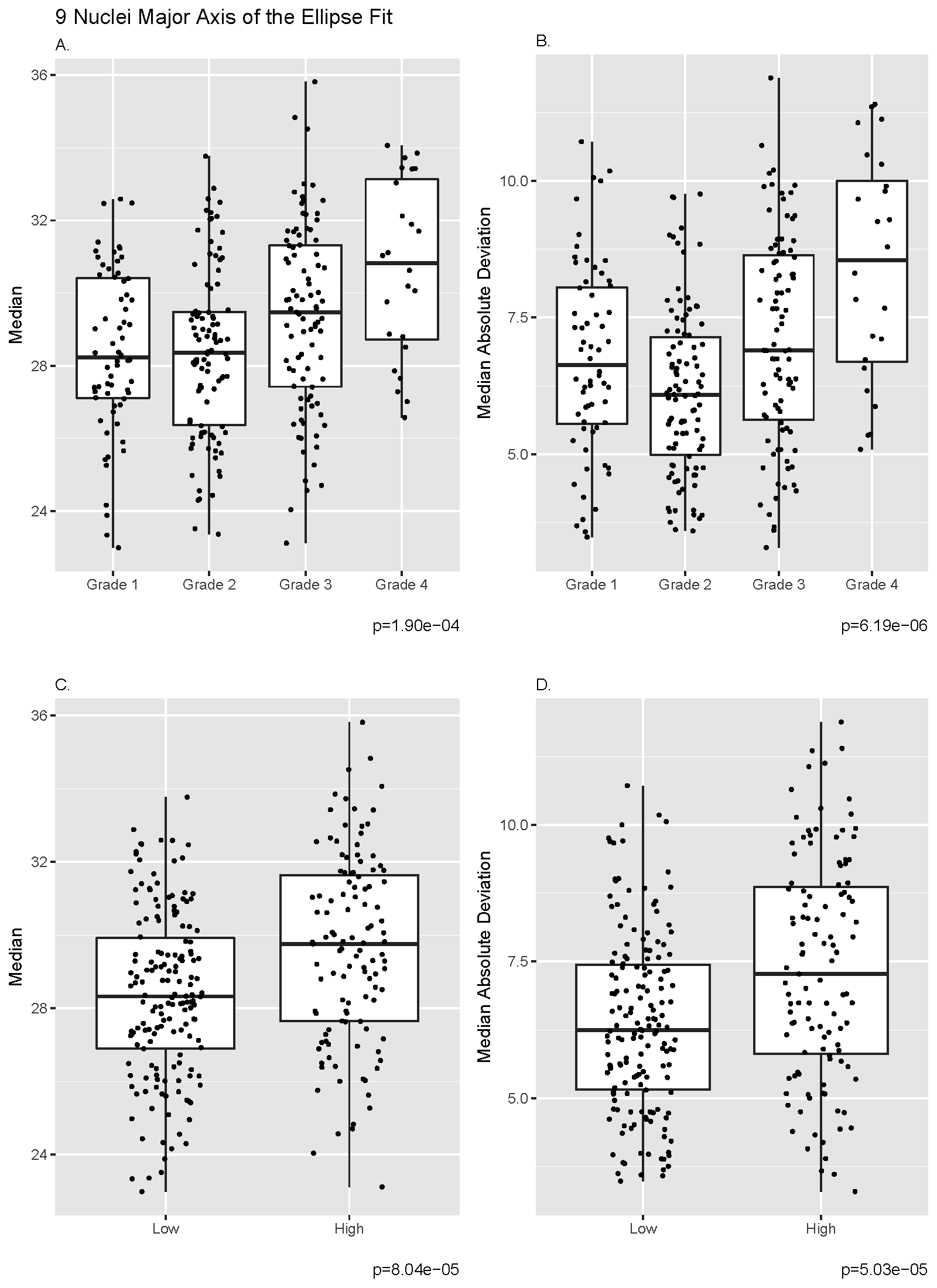

**Supplementary 5.** The association of the nine nuclear morphological features with Fuhrman’s grade in the 277 concordant cases (using the 2-tiered grading system).

|  | **Four-tiered Grading** | | **Two-tiered Grading** | |
| --- | --- | --- | --- | --- |
|  | **Median** | **MAD** | **Median** | **MAD** |
| **p-values** |  |  |  |  |
| Area | 9.20E-02 | **7.86E-04** | **3.36E-02** | **1.69E-04** |
| Roundness | **9.17E-06** | **6.43E-03** | **1.18E-05** | **6.63E-04** |
| Elongation | **5.78E-07** | **6.54E-06** | **1.30E-05** | **1.05E-02** |
| Flatness | **5.78E-07** | **6.54E-06** | **1.30E-05** | **1.05E-02** |
| Perimeter | **9.11E-06** | **8.10E-06** | **6.31E-06** | **1.08E-05** |
| Equivalent Spherical Perimeter | 9.20E-02 | **1.40E-04** | **3.36E-02** | **2.92E-05** |
| Equivalent Spherical Radius | 9.20E-02 | **1.40E-04** | **3.36E-02** | **2.92E-05** |
| Minor Axis of the Ellipse Fit | 7.21E-01 | **1.63E-04** | 3.87E-01 | **3.62E-05** |
| Major Axis of the Ellipse Fit | **1.90E-04** | **6.19E-06** | **8.04E-05** | **5.03E-05** |
| **FDR adjusted values** |  |  |  |  |
| Area | 1.03E-01 | **8.84E-04** | **3.78E-02** | **2.53E-04** |
| Roundness | **2.06E-05** | **6.43E-03** | **2.92E-05** | **8.52E-04** |
| Elongation | **2.60E-06** | **1.82E-05** | **2.92E-05** | **1.05E-02** |
| Flatness | **2.60E-06** | **1.82E-05** | **2.92E-05** | **1.05E-02** |
| Perimeter | **2.06E-05** | **1.82E-05** | **2.92E-05** | **8.15E-05** |
| Equivalent Spherical Perimeter | 1.03E-01 | **2.10E-04** | **3.78E-02** | **8.15E-05** |
| Equivalent Spherical Radius | 1.03E-01 | **2.10E-04** | **3.78E-02** | **8.15E-05** |
| Minor Axis of the Ellipse Fit | 7.21E-01 | **2.10E-04** | 3.87E-01 | **8.15E-05** |
| Major Axis of the Ellipse Fit | **3.42E-04** | **1.82E-05** | **1.45E-04** | **9.05E-05** |

Median absolute deviation, MAD; red text indicates a value that is p<0.05 or false discovery rate (fdr) <0.05.

**Supplementary 6.** Contingency tables of Fuhrman’s grade between TCGA and Pathologist 1 with Pathologist 2 among the 118 discordant cases.

|  | 4-tiered grading system | | | |  | 2-tiered grading system | |
| --- | --- | --- | --- | --- | --- | --- | --- |
|  | Pathologist 2 | | | |  | Pathologist 2 | |
|  | Grade 1 | Grade 2 | Grade 3 | Grade 4 |  | Low Grade | High Grade |
| TCGA |  |  |  |  | TCGA |  |  |
| Grade 1 | 0 | 0 | 0 | 0 | Low Grade | 9 | 19 |
| Grade 2 | 0 | 9 | 16 | 3 | High Grade | 53 | 37 |
| Grade 3 | 2 | 46 | 29 | 1 |  |  |  |
| Grade 4 | 0 | 5 | 6 | 1 |  |  |  |
| Pathologist 1 |  |  |  |  | Pathologist 1 |  |  |
| Grade 1 | 2 | 11 | 3 | 0 | Low Grade | 53 | 37 |
| Grade 2 | 0 | 40 | 32 | 2 | High Grade | 9 | 19 |
| Grade 3 | 0 | 9 | 15 | 3 |  |  |  |
| Grade 4 | 0 | 0 | 1 | 0 |  |  |  |

**Supplementary 7.** The association of re-assigned grades for discordant cases.

|  | **Re-assigned Consensus Manual Grade** | | |
| --- | --- | --- | --- |
|  | **Hazard**  **Ratio** | **(95% CI)** | ***p*-value** |
| **A.** **Most frequent grade among TCGA, Pathologist 1, and Pathologist 2** | | | |
| Model A: Crude | 1.09 | (0.63, 1.89) | 0.75 |
| Model B: Adjusted for Age and Gender | 1.21 | (0.70, 2.10) | 0.50 |
| Model C: Adjusted for Age, Gender, and Stage | 1.15 | (0.66, 2.00) | 0.62 |
| **B.** **Most frequent grade among Pathologist 1, Pathologist 2, and Computer** | | | |
| Model A: Crude | 1.33 | (0.77, 2.29) | 0.31 |
| Model B: Adjusted for Age and Gender | 1.42 | (0.81, 2.47) | 0.22 |
| Model C: Adjusted for Age, Gender, and Stage | 1.24 | (0.70, 2.20) | 0.46 |

Confidence Interval, CI
